# Gene model for the ortholog of *DENR* in *Drosophila grimshawi*

**DOI:** 10.64898/2026.07.14.738501

**Authors:** Megan E. Lawson, Kylee Sanow, Mihai Fratian, Madelyn Matura, Inayah Burton, Chinmay P. Rele, Jeffrey S. Thompson, Solomon Tin Chi Chak, Kellie S. O’Rourke

## Abstract

Gene model for the ortholog of *Density regulated protein* (*DENR*) in the May 2011 (Agencourt dgri_caf1/DgriCAF1) Genome Assembly (GenBank Accession: GCA_000005155.1) of *D. grimshawi*. This ortholog was characterized as part of a developing dataset to study the evolution of the Insulin/insulin-like growth factor signaling pathway (IIS) across the genus *Drosophila* using the Genomics Education Partnership gene annotation protocol for Course-based Undergraduate Research Experiences.

## Introduction

“Computational gene predictions in non-model organisms often can be improved by careful manual annotation and curation, allowing for more accurate analyses of gene and genome evolution (Mudge and Harrow 2016; Tello-Ruiz et al., 2019). The Genomics Education Partnership (thegep.org) uses web-based tools to allow undergraduates to participate in course-based research by generating manual annotations of genes in non-model species (Rele et al., 2023). These models of orthologous genes across species, such as the one presented here, then provide a reliable basis for further evolutionary genomic analyses when made available to the scientific community. The particular gene ortholog described here *Density regulated protein* (*DENR*) in *D. grimshawi* was characterized as part of a developing dataset to study the evolution of the Insulin/insulin-like growth factor signaling pathway (IIS) across the genus Drosophila.” (Myers et al., 2024).

“The IIS pathway is a highly conserved signaling pathway in animals and is central to mediating organismal responses to nutrients (Hietakangas and Cohen 2009; Grewal 2009)” (Myers et al., 2024). “*DENR* was first discovered in a human teratocarcinoma cell line because its concentration in cells increased with cell density (Deyo et al., 1998). Subsequent bioinformatic and biochemical analyses showed that the protein is conserved across eukaryotes and functions in non-canonical translation initiation (Fleischer et al., 2006; Skabkin et al., 2010). *D. melanogaster* flies homozygous for a null, knockout allele of the gene encoding *DENR* (FBgn0030802), die as pharate adults, showing a larval-like epidermis and reduced proliferation of histoblast cells (Schleich et al., 2014). Subsequent experiments using both RNAi in S2 cells and the knockout allele in larvae showed that DENR is required, along with its interacting partner MCT-1, for the proper expression regulation of a subset of transcripts required for cell cycle progression and growth. In particular, the loss of *DENR* reduces expression of the insulin receptor and makes larvae less sensitive to insulin signaling (Schleich et al., 2014), thus implicating DENR in the regulation of the insulin signaling pathway.” (Laskowski et al., 2024).

“D. grimshawi (NCBI:txid7222) is a member of the Picture Wing clade (sensu Kaneshiro et al., 1995) of the Hawaiian Drosophila. Molecular analyses place the monophyletic Hawaiian Drosophila as sister to Scaptomyza clade, which both are nested within the Drosophila subgenus of the genus Drosophila(Kambysellis et al., 1995; Baker and DeSalle 1997). The Picture Wings are so called due to their dramatically pigmented wings. D. grimshawi was first described by Oldenberg (1914), and is found in high elevation cool tropical rainforest on the Maui Complex islands where they breed on rotting vegetation (Carson et al., 1970; Carson 1983).” (Lawson et al., 2025).

We propose a gene model for the *D. grimshawi* ortholog of the *D. melanogaster Density regulated protein* (*DENR*) gene. The genomic region of the ortholog corresponds to the uncharacterized protein XP_001991685.1 (Locus ID LOC6564724) in the May 2011 (Agencourt dgri_caf1/DgriCAF1) Genome Assembly of *D. grimshawi* (GenBank Accession: GCA_000005155.1). This model is based on RNA-Seq data from *D. grimshawi* (Yang et al. 2018; SRP073087) and *DENR* in *D. melanogaster* using FlyBase release FB2024_02 (GCA_000001215.4; Gramates et al., 2022; Jenkins et al., 2022; Larkin et al., 2021).The Genomics Education Partnership maintains a mirror of the UCSC Genome Browser (Kent WJ et al., 2002; Gonzalez et al., 2021), which is available at https://gander.wustl.edu.

## Results

### Synteny

The target gene, *DENR*, occurs on chromosome X in *D. melanogaster* and is flanked upstream by *CG4880* and *CG13002* and downstream by *RNA polymerase III subunit I* (*Polr3I*) and *Nitrogen permease regulator-like 2* (*Nprl2*). The *tblastn* search of *D. melanogaster* DENR-PA (query) against the *D. grimshawi* (GenBank Accession: GCA_000005155.1) Genome Assembly (database) placed the putative ortholog of *DENR* within scaffold scaffold_15203 (CH916370.1) at locus LOC6564724 (XP_001991685.1)— with an E-value of 4e-63 and a percent identity of 52.72%. Furthermore, the putative ortholog is flanked upstream by LOC6564723 (XP_001991684.1) and LOC6564585 (XP_001991683.2), which correspond to *Nprl2* and *Axs* in *D. melanogaster* (E-value: 0.0 and 0.0; identity: 92.84% and 84.83%, respectively, as determined by *blastp*; **Figure 1A**, Altschul et al., 1990). The putative ortholog of *DENR* is flanked downstream by LOC6564725 (XP_001991686.1) and LOC6564726 (XP_001991687.1), which correspond to *Polr3I* and *CG4880* in *D. melanogaster* (E-value: 2e-67 and 3e-55; identity: 73.08% and 35.25%, respectively, as determined by *blastp*). The putative ortholog assignment for *DENR* in *D. grimshawi* is supported by the following evidence: synteny of the genomic neighborhood is partially conserved, and additionally while the locations of *Nprl2* and *CG4880* are not syntenic, they both remain present in the genomic neighborhood of *DENR*. All *BLAST* results used to determine orthology indicate very high-quality matches, as well.

**Figure 1:**
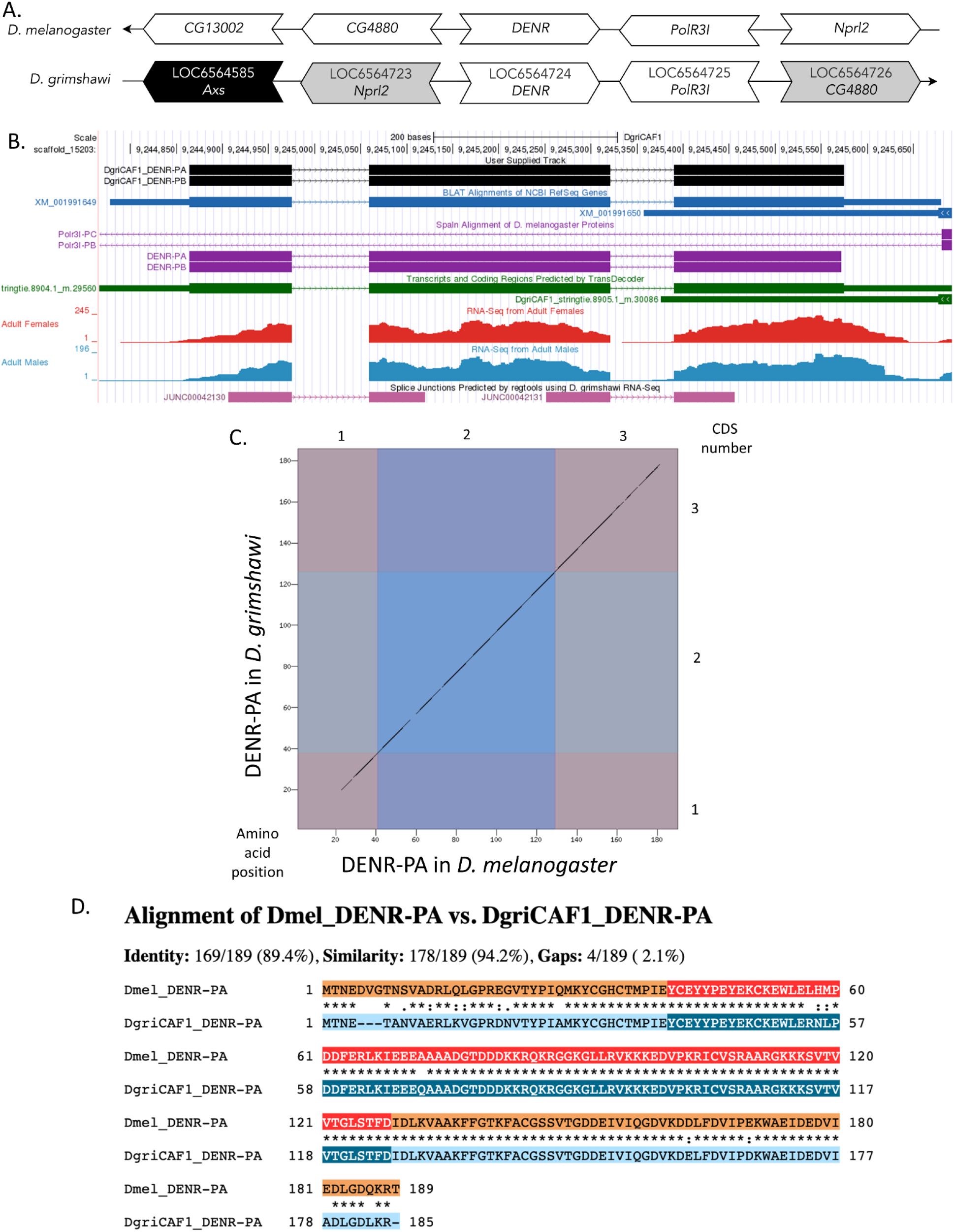
**(A) Synteny comparison of the genomic neighborhoods for *DENR* in *Drosophila melanogaster* and *D. grimshawi***. Thin underlying arrows indicate the DNA strand within which the target gene–*DENR*–is located in *D. melanogaster* (top) and *D. grimshawi* (bottom). The thin arrow pointing to the right indicates that *DENR* is on the positive (+) strand in *D. grimshawi*, and the thin arrow pointing to the left indicates that *DENR* is on the negative (-) strand in *D. melanogaster*. The wide gene arrows pointing in the same direction as *DENR* are on the same strand relative to the thin underlying arrows, while wide gene arrows pointing in the opposite direction of *DENR* are on the opposite strand relative to the thin underlying arrows. White gene arrows in *D. grimshawi* indicate orthology to the corresponding gene in *D. melanogaster*, black gene arrows indicate non-orthology, and gray gene arrows indicate that a gene is present in both neighborhoods but in different locations relative to the target gene. Gene symbols given in the *D. grimshawi* gene arrows indicate the orthologous gene in *D. melanogaster*, while the locus identifiers are specific to *D. grimshawi*. **(B) Gene Model in GEP UCSC Track Data Hub** (Raney et al. 2014). The coding-regions of *DENR* in *D. grimshawi* are displayed in the User Supplied Track (black); coding CDSs are depicted by thick rectangles and introns by thin lines with arrows indicating the direction of transcription. Subsequent evidence tracks include BLAT Alignments of NCBI RefSeq Genes (dark blue, alignment of Ref-Seq genes for *D. grimshawi*), Spaln of D. melanogaster Proteins (purple, alignment of Ref-Seq proteins from *D. melanogaster*), Transcripts and Coding Regions Predicted by TransDecoder (dark green), RNA-Seq from Adult Females and Adult Males (red and light blue, respectively; alignment of Illumina RNA-Seq reads from *D. grimshawi*), and Splice Junctions Predicted by regtools using *D. grimshawi* RNA-Seq (Yang et al. 2018; SRP073087). Splice junctions shown have a minimum read-depth of 10 with 100-499 supporting reads indicated in pink. **(C) Dot Plot of DENR-PA in *D. melanogaster* (*x*-axis) vs. the orthologous peptide in *D. grimshawi* (*y*-axis)**. Amino acid number is indicated along the left and bottom; CDS number is indicated along the top and right, and CDSs are also highlighted with alternating colors. Gaps in the dot plot result from decreased sequence similarity. **(D) Protein alignment of DENR-PA in *D. melanogaster* (top row) vs. the orthologous peptide in *D. grimshawi* (bottom row)**. The alternating colored rectangles represent adjacent CDSs. The symbols in the match line denote the level of similarity between the aligned residues. An asterisk (*) indicates that the aligned residues are identical. A colon (:) indicates the aligned residues have highly similar chemical properties—roughly equivalent to scoring > 0.5 in the Gonnet PAM 250 matrix (Gonnet et al., 1992). A period (.) indicates that the aligned residues have weakly similar chemically properties—roughly equivalent to scoring > 0 and ≤ 0.5 in the Gonnet PAM 250 matrix. A space indicates a gap or mismatch when the aligned residues have a complete lack of similarity—roughly equivalent to scoring ≤ 0 in the Gonnet PAM 250 matrix. This protein alignment shows that while there is decreased sequence similarity within the beginning portion of the first CDS and the end of the last CDS of DENR-PA in *D. melanogaster* and *D. grimshawi*, there is still high conservation of the peptide sequence and the chemical properties of these regions of the peptide.

### Protein Model

*DENR* in *D. grimshawi* has two protein-coding isoforms (DENR-PA and DENR-PB; **Figure 1B**). Isoforms DENR-PA and DENR-PB are identical and contain three protein-coding exons. Relative to the ortholog in *D. melanogaster*, the CDS number and protein isoform count are conserved, as *DENR-RA* and *DENR-RB* are also identical with three coding CDSs in *D. melanogaster*. The sequence of DENR-PA in *D. grimshawi* has 89.4% identity with the protein-coding isoform DENR-PA in *D. melanogaster*, as determined by *blastp* (**Figure 1C**). Small regions of low sequence identity were found in the first and last CDSs, but the protein alignment (**Figure 1D**) indicates that these regions still have very high chemical similarity across the two species. Coordinates of this curated gene model (*DENR-RA* and *DENR-RB*) are stored by NCBI at GenBank/BankIt (accession BK065241 and BK065242, respectively). These data are also archived in the CaltechDATA repository (see “Extended Data” section below).

## Methods

Detailed methods including algorithms, database versions, and citations for the complete annotation process can be found in Rele et al. (2023).

## Acknowledgements

We would like to thank Wilson Leung for developing and maintaining the technological infrastructure that was used to create this gene model and Laura K. Reed for overseeing the project.

## Supplemental Files

1. Zip file containing FASTA, PEP, GFF files for the gene model

## Metadata

Bioinformatics, Genomics, *Drosophila*, Genotype Data, New Finding

